# The oscillation of mitotic kinase governs cell cycle latches

**DOI:** 10.1101/2021.12.07.471646

**Authors:** Bela Novak, John J Tyson

## Abstract

In order to transmit a eukaryotic cell’s genome accurately from mother cell to daughter cells, it is essential that the basic events of the cell division cycle (DNA synthesis and mitosis) occur once and only once per cycle, i.e., that a cell progresses irreversibly from G_1_ to S to G_2_ to M and back to G_1_. Irreversible progression through the cell cycle is assured by a sequence of ‘latching’ molecular switches, based on molecular interactions among cyclin-dependent kinases and their auxiliary partners. Positive feedback loops (++ or −−) create bistable switches with latching properties, and negative feedback loops drive progression from one stage to the next. In budding yeast (*Saccharomyces cerevisiae*) these events are coordinated by double-negative feedback loops between Clb-dependent kinases (Clb1-6) and their antagonists (APC:Cdh1 and Sic1). If the coordinating signal is compromised, either by deletion of Clb1-5 proteins or expression of non-degradable Clb2, then irreversibility is lost and yeast cells exhibit multiple rounds of DNA replication or mitotic exit events (Cdc14 endocycles). Using mathematical modelling of a stripped-down control network, we show how endocycles arise because the switches fail to latch, and the gates swing back and forth by the action of the negative feedback loops.

## Introduction

The eukaryotic cell division cycle is typically divided into four phases: G_1_, S, G_2_ and M, where S = DNA synthesis, M = mitosis and cell division, G_1_ = first gap (unreplicated chromosomes), and G_2_ = second gap (replicated chromosomes). In 1996, Kim Nasmyth [1] proposed that, rather than four transient phases, the cell cycle should be thought of as two self-maintaining states: G_1_ and S-G_2_-M. The transition from G_1_ to S-G_2_-M (called the ‘Restriction Point’ in mammalian cells and ‘Start’ in yeast) is a major decision point in the reproductive life of a cell, when the cell commits itself to a new round of DNA replication and mitosis. The reverse transition, from S-G_2_-M back to G_1_, is the second major decision point, when the cell, after confirming that all replicated chromosomes are properly aligned on the mitotic spindle, activates the cell-division process (anaphase, telophase, cell division). This decision point is called the ‘Spindle Assembly Checkpoint’ [2]. In Nasmyth’s view, cell proliferation is an alternating sequence of transitions from G_1_ to S-G_2_-M and back again, as a cell lineage flips between these two alternative cell-cycle states.

Furthermore, Nasmyth correlated these two states (in budding yeast) to the activity of a family of cyclin-dependent kinases (CDKs) that trigger S-G_2_-M events (Cdk1:Clb5-6, which trigger DNA synthesis, and Cdk1:Clb1-4, which trigger chromosome condensation and chromosome alignment on the mitotic spindle) by phosphorylating specific proteins involved in these events. Consequently, the S-G_2_-M state is characterized by CDK activity rising to a high level and significant phosphorylation of S- and M phase-related proteins. Conversely, G_1_ phase of the cell cycle is characterized by low CDK activity and CDK-targeted proteins mostly dephosphorylated. The two states are ‘self-maintaining’, according to Nasmyth, because of the interactions of CDKs with E3 ubiquitin-ligases (UbLs) that label cyclin molecules for degradation by proteasomes, and with stoichiometric inhibitors (CKIs) that bind tightly to CDKs and competitively inhibit the phosphorylation of other substrates. Because CDKs phosphorylate both UbLs and CKIs (inhibiting UbL activity and priming CKIs for ubiquitination and degradation), CDKs and UbLs/CKIs are <u>mutually</u> antagonistic. Intuitively, Nasmyth understood (see Figure 3 of his paper) that these mutually antagonistic interactions could create two self-maintaining states of the CDK-control network and, thus, of cell cycle physiology.

At about the same time, we proposed similar ideas from a modeler’s perspective [3, 4]. Our idea was to associate these self-maintaining states of the cell cycle with alternative, stable steady-state solutions of the biochemical kinetic equations that describe the time evolution of the CDK-control network. In this view, G_1_ is a stable steady state of high activity of the CDK antagonists and low CDK activity, and S-G_2_-M is a state of inactive antagonists, enabling CDK activity to rise to a stable steady state of high activity promoting entry into mitosis. This ‘bistable’ control system is illustrated schematically by the vertical line in the centre of Figure 1. At the top of the line is a stable G_1_ state; the arrows pointing toward • indicate that small perturbations away from the steady state return directly to G_1_. At the bottom of the line is a stable M state (the end state of the S-G_2_-M sequence). The two stable steady states (•) are separated by an unstable steady state (○), which is an inevitable correlate of bistability even though it is experimentally unreachable. The line is oriented with high activity of antagonists at the top and high activity of CDKs at the bottom. For a cell to leave the G_1_ state and enter the S-G_2_-M sequence, it must activate a transcription factor (TF) that promotes the synthesis of a ‘G_1_ cyclin’ that can overwhelm the antagonists of the ‘S-G_2_-M cyclins’. The G_1_ cyclin-dependent kinase destabilizes the G_1_ state by forcing it to coalesce with the unstable steady state; as indicated in Figure 1 by the red line (the locus of stable G_1_ states) coalescing with the grey dashed line (the locus of unstable steady states) as TF increases. In the parlance of dynamical systems theory, the coalescence point is called a ‘saddle-node’ bifurcation. Beyond the bifurcation point, the only stable attractor of the dynamical system is the S-G_2_-M sequence (the green line at the bottom of Figure 1). As a consequence of entering this sequence, the TF that initiated the transition is inactivated, and subsequent progression through the division cycle brings the cell into mitosis (M, a stable steady state). To exit mitosis, divide and return the daughter cells to G_1_, the mitotic cell must activate some ‘exit proteins’ (EP) that promote anaphase (sister chromatid separation) and degradation of mitotic cyclins, so that the CDK antagonists can reassert themselves. This transition is illustrated to the left of centre in Figure 1, where the locus of S-G_2_-M states (the green line) coalesces with the unstable steady state at a saddle-node bifurcation induced by sufficiently high activity of EP. Beyond this point, the only stable attractor of the dynamical system is the G_1_ state (the red line at the top of the figure). As before, once EP has induced the transition, it is no longer needed; negative feedback interactions remove EP, and the daughter cells are left in the stable G_1_ state, with neither active EP nor active TF.

**Figure 1.**
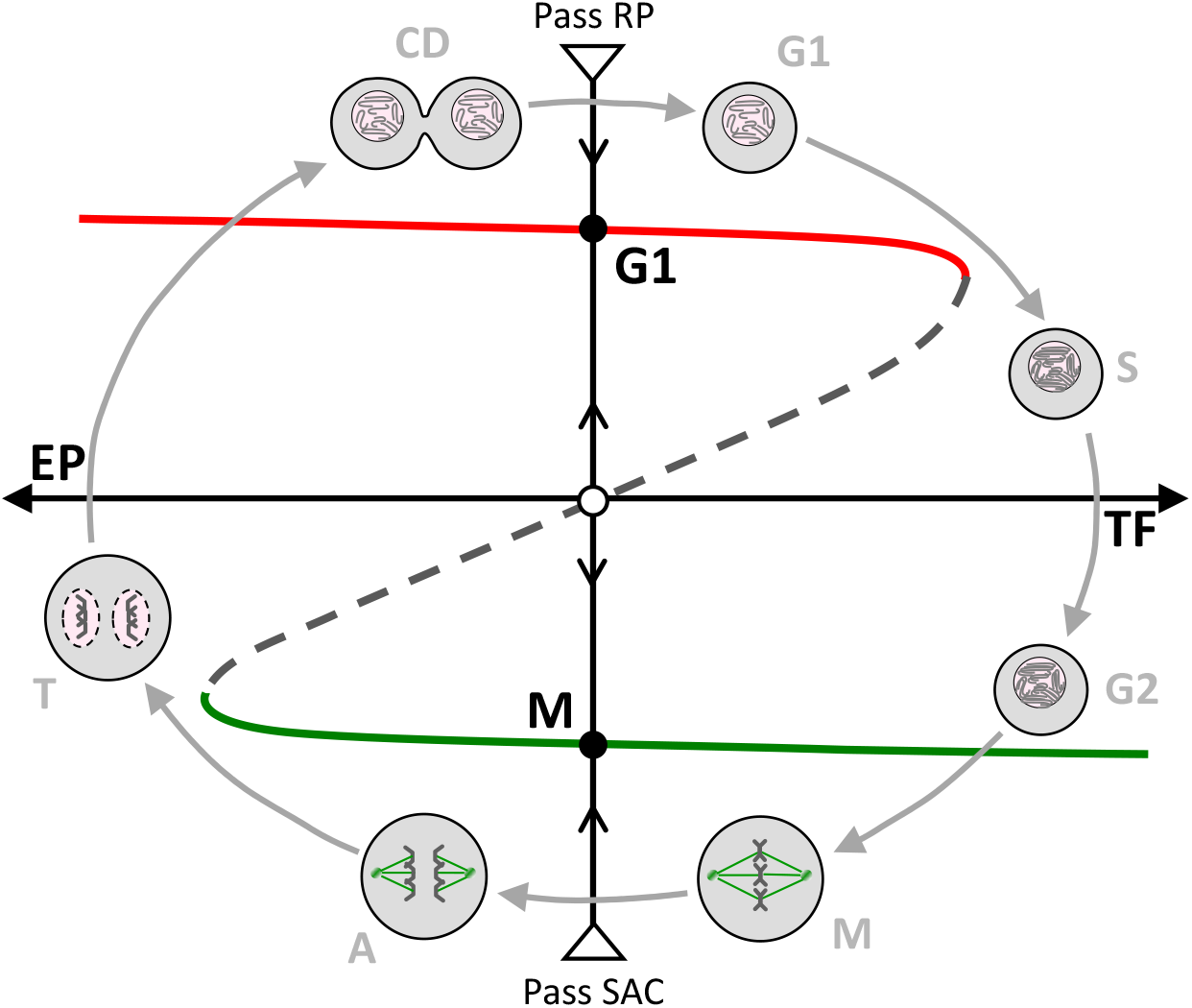
The eukaryotic cell division cycle viewed as a latching switch flipped by promoters (TF and EP) of the G_1_-S transition and the M-G_1_ transition (respectively). Around the outside we display schematic images of a cell progressing from G_1_ to S to G_2_ to metaphase (M). Within the cycle we indicate that the interactions of CDKs and their antagonists (UbLs and CKIs) create a bistable control system (central vertical line) with a stable G_1_ state (CDK inactive, UbL and CKI active) and a stable M state (CDK active, UbL and CKI inactive), separated by an unstable state (○). To flip the switch from G_1_ to M, the cell must pass the Restriction Point (RP) and activate a transcription factor (TF) that induces synthesis of proteins that inhibit UbL and CKI, giving CDK a chance to establish its dominance. After the transition is made (flipping from the red state to the green state), the TF is inactivated and its encoded proteins degraded, and active CDK maintains the switch in the stable M state. To exit mitosis (metaphase to anaphase to telophase to G_1_), the cell must satisfy the Spindle Assembly Checkpoint (SAC) and activate a set of exit proteins (EP) that depress CDK activity so that UbL and CKI can re-establish their dominance. After the transition is complete, the EPs are removed and the antagonists maintain the switch in the stable G_1_ state.

At the stable G_1_ state—the Restriction Point (RP) in mammals—the cell is waiting for conditions to be conducive to a new round of DNA replication and division. These conditions include stimulation by growth factors, absence of DNA damage, and (to some extent) sufficiently large size. At the S-G_2_-M state, the cell is waiting at the Spindle Assembly Checkpoint (SAC) for notification that all replicated chromosomes are properly aligned on the mitotic spindle [2].

(In many cell types the G_2_-M transition is guarded by a third checkpoint that blocks entry into M phase until all chromosomes are fully replicated and any DNA damage suffered since the G_1_-S transition has been repaired. This checkpoint is absent in budding yeast cells (the focus of our study); so we will not consider it in what follows. Nonetheless, the G_2_-M checkpoint can be modelled by exactly the same methods, as described in other publications [5-7].)

Following Nasmyth’s lead, the dynamical picture in Figure 1 views cell proliferation as a series of flips of a bistable switch back-and-forth between G_1_ (a cell with unreplicated DNA) and S-G_2_-M (a cell with replicated DNA). The switch functions like an old-fashioned mechanical light switch (see Supplemental Figure 1). The lever must be pushed a sufficient distance in one direction (by TF) to flip from OFF to ON (from G_1_ to S-G_2_-M), after which the switch latches into the ON position. To turn the ‘light’ off, the lever must be pushed sufficiently far in the opposite direction (by EP), after which it latches OFF. The latches lock in place because of the antagonistic interactions between S-G_2_-M cyclins and their ‘enemies,’ the UbLs and CKIs that oppose S phase-promoting and M phase-promoting CDKs. <u>For the switches to latch properly, the S-G_2_-M cyclins must oscillate between low activity (G_1_) and high activity (S-G_2_-M) states, and the inducers (TF and EP) must be under negative-feedback control</u> (i.e., TF must be inactivated after the cell enters S-G_2_-M, and EP must be inactivated after the cell returns to G_1_).

We have described this concept of cell cycle control at length in previous publications [8, 9], and our theoretical ideas have been verified by elegant experiments with budding yeast: the irreversible characteristics of Start and mitotic exit were confirmed by Cross et al. [10] and by Lopez-Aviles et al. [11], respectively. In this paper we want to consider some consequences (for cell cycle progression) of perturbing the underlying oscillation of mitotic cyclins by mutation. It is known that, in many cell types, if mitotic cyclins are deleted (so the cells cannot enter mitosis), then the negative feedback control on TF generates oscillations in S phase-promoting cyclins, i.e., repeated rounds of DNA replication without mitosis (called ‘endoreplication’ [12]). On the other hand, if non-degradable mitotic cyclins are expressed in budding yeast cells so that total mitotic CDK activity cannot fall to a low level on exit from mitosis, then the cells exhibit sustained oscillations of an EP (namely, Cdc14 phosphatase; hence, ‘Cdc14 endocycles’ [13, 14]). In the following sections we show how these perturbations of mitotic CDK activity change the picture in Figure 1 to generate these endocycles.

### A simple model of the budding yeast cell cycle

To illustrate our view of endocycles, we use a simple model of the budding yeast cell cycle (Figure 2A), constructed along the lines of the comprehensive model by Chen et al. [15]. At the heart of the network are double-negative interactions between ClbS and ClbM (Cdk1:Clb5-6 and Cdk1:Clb1-4, respectively) and their inhibitory substrates: Cdh1 (the APC:APC complex, a UbL) and Sic1 (a CKI). Cdh1 is inactivated by phosphorylation by ClbS and ClbM-kinases [16-18], while Sic1 is targeted to SCF-dependent proteolytic degradation by the same kinases [19]. Cln denotes the G_1_-cyclin dependent kinases Cdk1:Cln1-2, which can phosphorylate and incapacitate both Cdh1 and Sic1 [20], but, unlike ClbS and ClbM, Cln-kinase is not destroyed by Cdh1 or inhibited by Sic1. Cln synthesis is upregulated by its transcription factor, SBF. SBF regulation is complex: it is activated by a transcriptional positive feedback loop [21] and it is inhibited by ClbM-kinase activity [22], i.e., by the pathway SBF → Cln –I Cdh1 –I ClbM –I SBF. Finally, note that the level of ClbS is regulated by two negative feedback loops involving its transcription factor, MBF; namely MBF → Nrm1 –I MBF and MBF → ClbS –I Cdh1 –I Nrm1 –I MBF.

**Figure 2.**
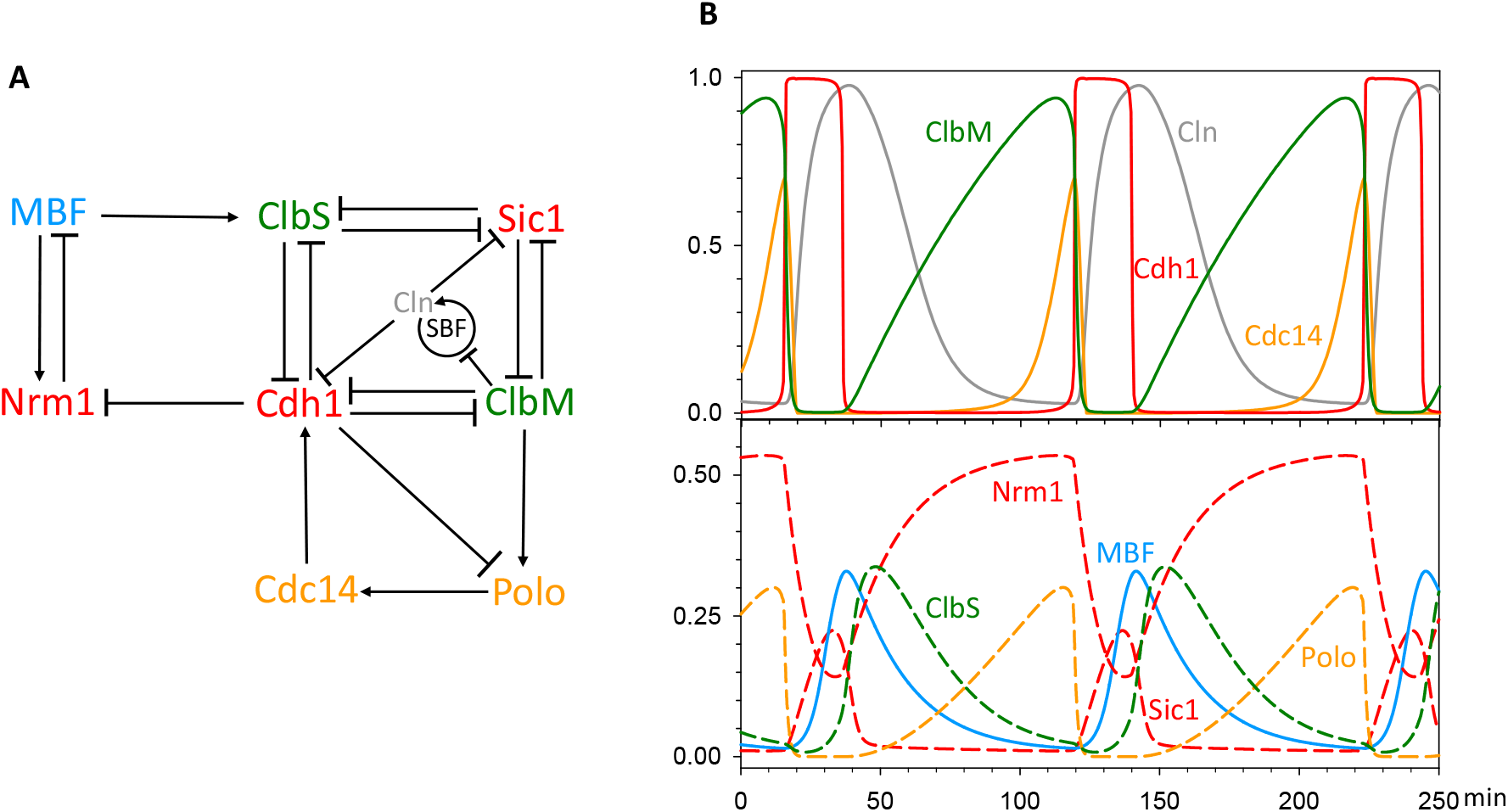
A simplified model of the budding yeast cell-cycle control system. **(A)** The influence diagram. Barbed arrows indicate positive influences (activation or synthesis). Blunt connectors indicate negative influences (inhibition or degradation). ClbS, ClbM and Cln are CDKs, Cdh1 is a UbL, Sic1 is a CKI, SBF and MBF are TFs, Polo (a kinase) and Cdc14 (a phosphatase) are EPs, Nrm1 is a transcriptional co-repressor. See text for further explanation. **(B)** Simulated time course of a mother cell undergoing three divisions. Upper panel. ClbM and Cdh1 are mutually antagonistic proteins, creating alternative stable states: G_1_ (Cdh1 active and ClbM falling rapidly), and S-G_2_-M (Cdh1 inactive and ClbM rising slowly). By upregulating ClbS and Cln, MBF and SBF push the cell from G_1_ into S-G_2_-M. Polo and Cdc14 induce the reverse transition. Lower panel. Behaviour of the other components of the wiring diagram. The time-courses are computed by numerical simulation of the nonlinear ODEs in Suppl. Text A

The first event of the Start transition is the activation of SBF and MBF, which drive the production of Cln and ClbS, respectively. As noted above, SBF and MBF are both under negative regulation, but by different pathways. We might consider our generic TF (in Figure 1) to be some combination of SBF and MBF. We shall focus on the role of MBF in the budding yeast cell cycle for reasons that will be apparent soon.

To simplify the mathematical description, we model—not wild-type budding yeast cells but—the cell cycle of *cdc20Δ pds1Δ clb5Δ* triple-mutant cells, which are viable [23]. Securin (Pds1) and S-phase cyclin (Clb5) are inhibitors of separase and Cdh1, respectively, and—in wild-type cells—they are removed by APC:Cdc20-dependent proteolysis during mitotic progression. In the absence of the inhibitors (Pds1 and Clb5) of cell cycle progression, the Cdc20 cell cycle activator is dispensable. In the triple mutant, the degradation of mitotic cyclin (ClbM) is dependent solely on APC:Cdh1, which is controlled by a negative feedback loop: ClbM → Polo → Cdc14 → Cdh1 –I ClbM. In this loop, Cdc14 is our generic EP, i.e., a mitotic exit phosphatase [24].

In Suppl. Text A we provide a set of ordinary differential equations (ODEs) that describe this simplified model of the budding yeast cell cycle. Numerical integration of the ODEs (for the parameter values given in Suppl. Text A) provides a reasonable semi-quantitative description (see Figure 2B) of the observed fluctuations of cell cycle regulators in budding yeast cells growing in in rich (glucose) medium, with a 100 min cycle time. Note that we do not include the size control mechanism that is operative at Start in budding-yeast daughter cells; therefore, these oscillations correspond to the cell division cycles of mother cells of the triple-mutant cell line.

Now we are positioned to convert our schematic diagram of bistability and hysteresis (Figure 1) into a precise bifurcation diagram (Figure 3a) calculated for the dynamical system derived from the wiring diagram in Figure 2A. To this end, we use Cdh1 as the dynamical variable and MBF and Cdc14 as the bifurcation parameters (TF and EP). (In Suppl. Figure 2 we draw the same bifurcation diagram using ClbM as the dynamical variable.) The Z-shaped curve in Figure 3A is the locus of steady-state activity of Cdh1 as a function of fixed MBF (for Cdc14 = 0) to the right, and as a function of fixed Cdc14 (for MBF = 0) to the left. It is important to recognize that each half of the Z-shaped curve is itself Z-shaped in the following sense. For Cdh1 as a function of MBF (to the right), the middle branch of the red curve connects to the lower branch at a <u>negative</u> value of MBF. If the ‘elbow’ of the Z on the upper branch of the red curve is the ‘upper threshold’ for the abrupt inactivation of Cdh1 as MBF increases, the elbow of the Z on the lower branch is the ‘lower threshold’ for abrupt activation of Cdh1 as MBF decreases. Because the lower threshold is at negative activity of MBF, Cdh1 cannot be re-activated as MBF activity falls to zero. This effect is crucial to the ‘latching’ behavior of the control system. As the ‘gate swings open’ and the cell transits from G_1_ to S-G_2_-M, the ‘spring (the negative feedback on MBF) pulls the gate closed again and the latch (the stability of the metaphase steady state)’ holds the cell in mitosis. Exactly the same latching mechanism holds on the left side of the diagram. There is a ‘lower threshold’ for Cdh1 inactivation as Cdc14 activity falls, but it occurs at <u>negative</u> value of Cdc14 activity. So the latch catches the gate at the G_1_ steady state.

**Figure 3.**
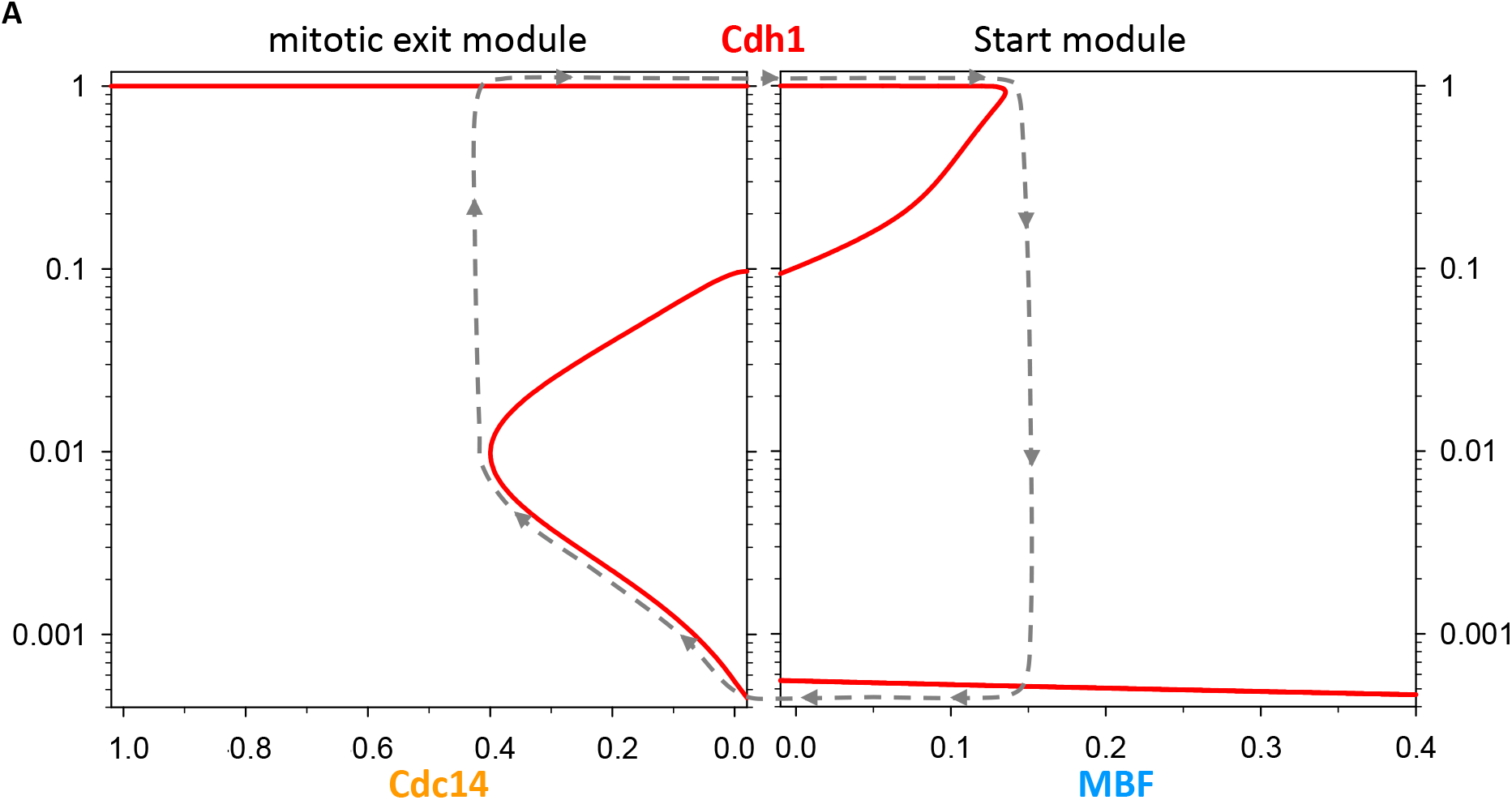
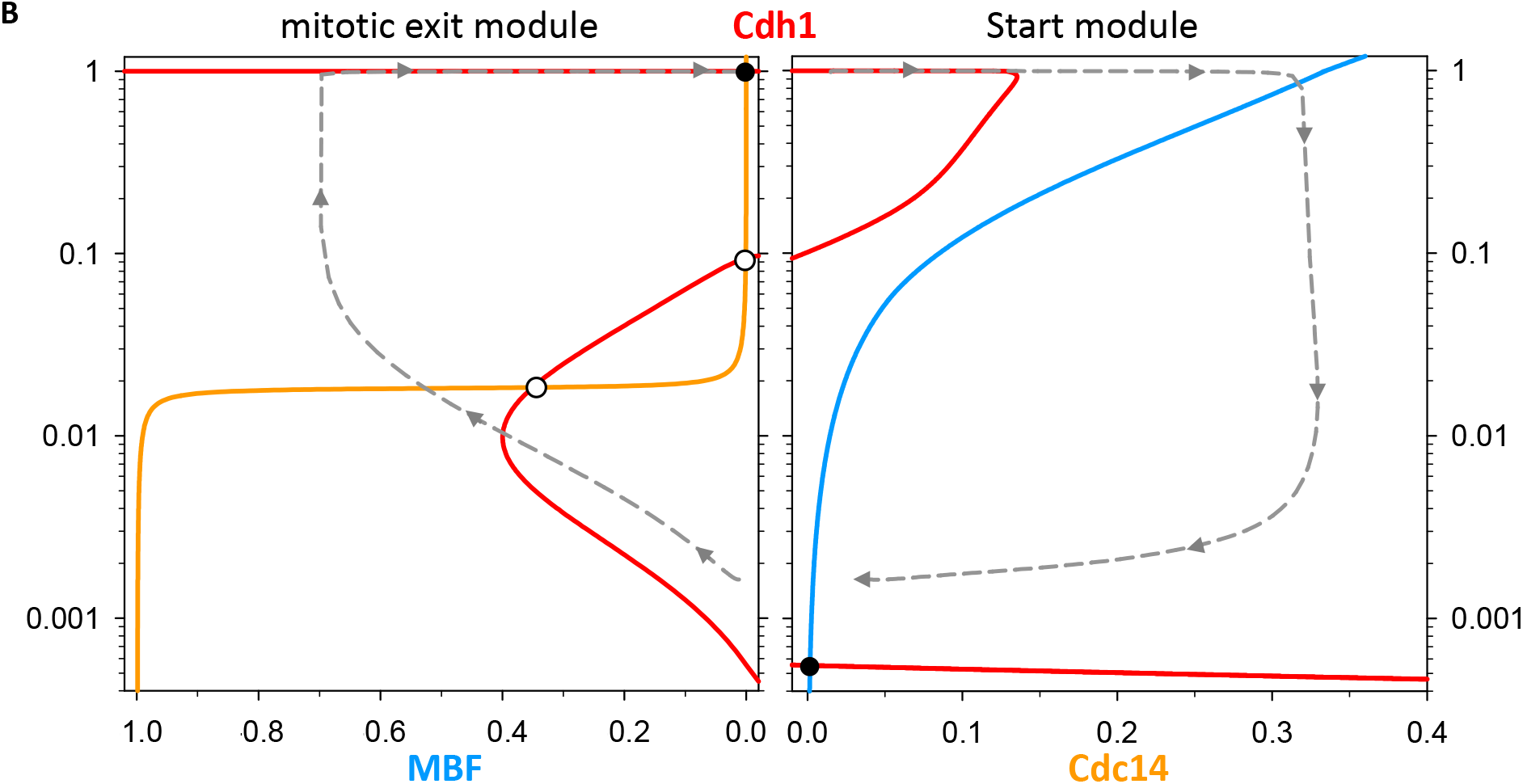
The budding yeast cell division cycle viewed as a latching switch, based on bifurcation curves computed from the differential equations representing the temporal dynamics of the CDK control system in Figure 2A. **(A)** On the right (left) we plot the steady state activity of Cdh1 as a function of fixed MBF activity (Cdc14 activity) for Cdc14 = 0 (MBF = 0). The grey dashed curve is a schematic diagram of the expected hysteresis loop. (**B)** Dynamical evolution of the control system. Red curve, the Cdh1 balance curve, as in panel a; blue curve, the MBF balance curve (steady-state value of MBF as a function of Cdh1, for Cdc14 = 0); orange curve, the Cdc14 balance curve (*mutatis mutandis*). Intersection points are steady states (•, stable; ○, unstable). Dashed line is the simulated trajectory in Figure 2B projected onto the bifurcation diagrams.

To gain more insight into cell cycle progression, we superimpose on the bifurcation diagram (see Figure 3B) the influence of Cdh1 activity on the steady-state levels of MBF and Cdc14. MBF becomes more active as Cdh1 activity increases (the blue curve), because Cdh1 degrades the MBF inhibitor, Nrm1 [25]. In contrast, Cdc14 activity decreases with increasing Cdh1 (the orange curve), because Cdh1 degrades the Cdc14 inhibitor, Polo [26].

We may think of the curves on Figure 3B as ‘pseudo-nullclines’: on the red curves, Cdh1 is at a pseudo-steady state (d[Cdh1]/dt ≈ 0), on the blue curve, d[MBF]/dt ≈ 0, and on the orange curve, d[Cdc14]/dt ≈ 0. We say ‘≈ 0’ because the actual rates of change depend on what the other variables of the dynamical system are doing at any particular time in a simulation. Wherever two pseudo-nullclines intersect is a potential steady state of the full set of differential equations; the black dot (•) on the left, at Cdh1 = 1, is a true steady state if MBF = 0, and the black dot on the right, at Cdh1 ≈ 0.0005, is a true steady state if Cdc14 = 0. By projecting onto Figure 3B the trajectory (dashed curves) of the dynamical system from the simulation in Figure 2B, we see that the pseudo-nullclines are useful in understanding how the dynamical system evolves in time. Nonetheless, at low Cdh1 activity the trajectories deviate considerably from the pseudo-nullclines, as should be expected, because numerical simulation of the full set of dynamical equations more accurately predicts the time-evolution of the control system than forecasts based on bifurcation curves derived from steady-state assumptions.

### Endoreplication cycle in the absence of mitotic cyclins

In fission yeast cells, shutting off the synthesis mitotic cyclin (Cdc13) leads to periodic rounds of DNA synthesis in the absence of mitosis [27]. These endoreplication cycles are driven by periodic expression of fission yeast’s S-phase cyclin (Cig2). In budding yeast, deletion of the four mitotic cyclins (Clb1-4) causes a G_2_ block because of persisting activity of the S-phase cyclin (Clb5) [28]. However, deletion of the *CLB5* gene supports periodic endoreplication cycles in the *clb1-5Δ* strain [29], which must be driven by an oscillation of the other S-phase cyclin (Clb6) in budding yeast. Interestingly, Clb6 is the only B-type cyclin that is targeted for degradation by an SCF-dependent mechanism [30], which evidently allows replication origin relicensing in the absence of Clbs 1-5.

We simulate the *clb1-5Δ* strain by reducing the synthesis of mitotic cyclins (ClbM) to zero (see Figure 4A). In this case, the cell cycle control network exhibits limit-cycle oscillations of ClbS activity driven by periodic synthesis and degradation. Cln-kinase activity is maintained at a constantly high level because SBF is constitutively active in the absence of inhibition by mitotic cyclins (Clb1-4) [22].

**Figure 4.**
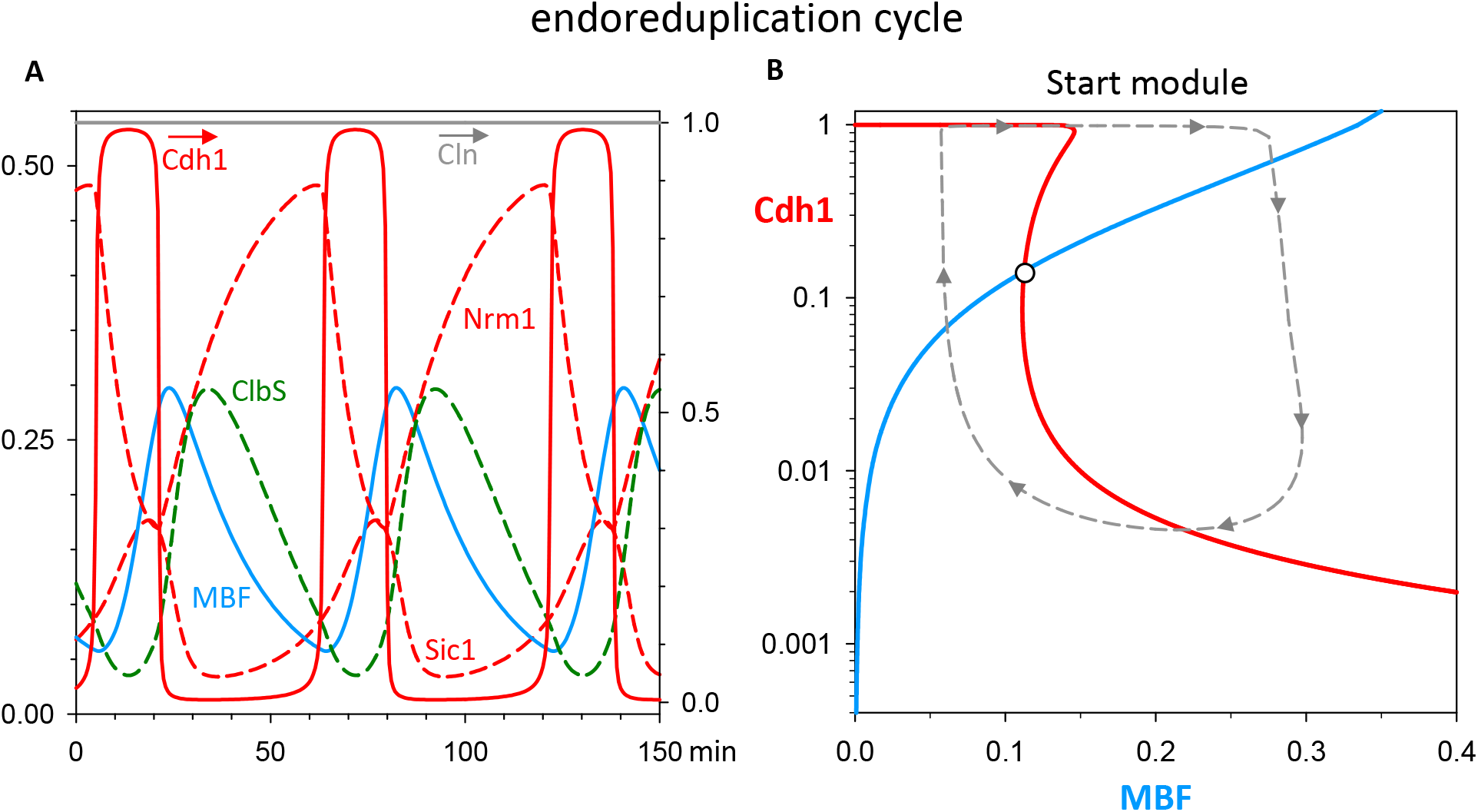
Endoreplication cycles in the *clb1-5Δ* mutant strain. **(A)** Numerical simulation of the model ODEs for *k*_*sclbm*_*’* = *k*_*sclbm*_ = 0, i.e., no synthesis of ClbM. Note the periodic expression of ClbS, driving periodic rounds of DNA replication; also, because ClbM is missing, SBF is constitutively active and Cln is constantly present at high level. **(B)** Pseudo-phase plane. Red curve is steady state activity of Cdh1 as a function of MBF (MBF drives expression of ClbS, which phosphorylates and inactivates Cdh1). Blue curve is steady state activity of MBF as a function of Cdh1 (Cdh1 degrades Nrm1, which is an inhibitor of MBF). Grey dashed line is the limit cycle in panel A, projected onto the MBF-Cdh1 plane. It is a ‘pseudo’ phase plane because the dynamical system has six variables, and the red and blue curves are not ‘nullclines.’ Nonetheless, the pseudo-phase plane gives a good idea of how endoreplication cycles arise in the model in the absence of ClbM.

Removing the synthesis of mitotic cyclins in the model causes a major change in the dynamics of the Start module (see Figure 4B). In the absence of mitotic cyclins, the irreversible (latching) nature of the Start transition is lost because the lower threshold of the Z-shaped Cdh1 balance curve, as a function of MBF, moves to a positive value of MBF activity. This change makes the steady state at the intersection of the two balance curves unstable, and the control system executes **limit cycle** oscillations indicated by the trajectory (dashed curve) from the numerical simulation. In Suppl. Figure S3 we show that these limit cycle oscillations arise by a SNIC bifurcation (saddle-node on an invariant circle).

The persistence of Cdh1 bistability is a consequence of our assumption that the S-phase cyclin (Clb6), in addition to its degradation by SCF, is also degraded by APC:Cdh1 during G_1_ phase. This assumption maintains the antagonistic relationship between Cdk1 and Cdh1 during endoreplication cycles. In case this assumption does not hold, the Cdh1 balance curve loses its S-shaped characteristic and becomes sigmoidal; nonetheless, the network still executes limit-cycle oscillations driven by time-delayed negative feedback alone.

### Cdc14 endocycles

The lack of mitotic cyclins converts the latching mechanism of the Start module into an autonomous oscillatory network which drives periodic DNA replication in the absence of mitosis. The opposite situation, when mitotic cyclin activity stays high, can be achieved in budding yeast cells by introducing the *CLB2kdΔ* gene (KEN and D-box deleted) encoding a functional, non-degradable Clb2 protein (ndClbM). The degradation of mitotic cyclins is a necessary requirement for dephosphorylation of mitotic substrates by counter-acting phosphatases. In budding yeast, the bulk of mitotic dephosphorylation is catalysed by Cdc14 phosphatase, which is transiently released from the nucleolus during mitotic exit [31]. In the presence of supra-threshold activity of non-degradable Clb2, exit from mitosis is blocked and Cdc14 release becomes periodic [13, 14].

If we express a constant low level of non-degradable ClbM in our model (e.g., ndClbM = 0.1 in Figure 5A), cell cycle progression is hardly perturbed and mitotic exit is not compromised. This can be confirmed by the large post-mitotic peak of Cln level, whose synthesis by SBF is very sensitive to inhibition by ClbM. Hence, ClbM activity must be very low as these cells exit mitosis, presumably because the non-degradable fraction of ClbM is inhibited by the peak of Sic1 in early G_1_. Low ClbM activity would allow re-licensing of chromosome replication origins and a normal cell cycle. At a higher level of ndClbM, the cell is unable to leave mitosis and start a new cell division cycle. For example, at ndClbM = 0.4 (Figure 5B), total ClbM > 0.4 and Sic1 < 0.02, so there is sufficient ClbM activity at all times to block re-licensing of replication origins; hence, the genome cannot be replicated even though ClbS is oscillating. Nonetheless, Cdc14 is periodically inactivated and activated (i.e., sequestered in and released from nucleoli) by the negative feedback loop Polo → Cdc14 → Cdh1 –|Polo. For sufficiently large amounts of non-degradable ClbM (ndClbM > 0.575), Cdc14 executes small amplitude oscillations around a high background level indicating that the phosphatase is not sequestered in the nucleolus.

**Figure 5.**
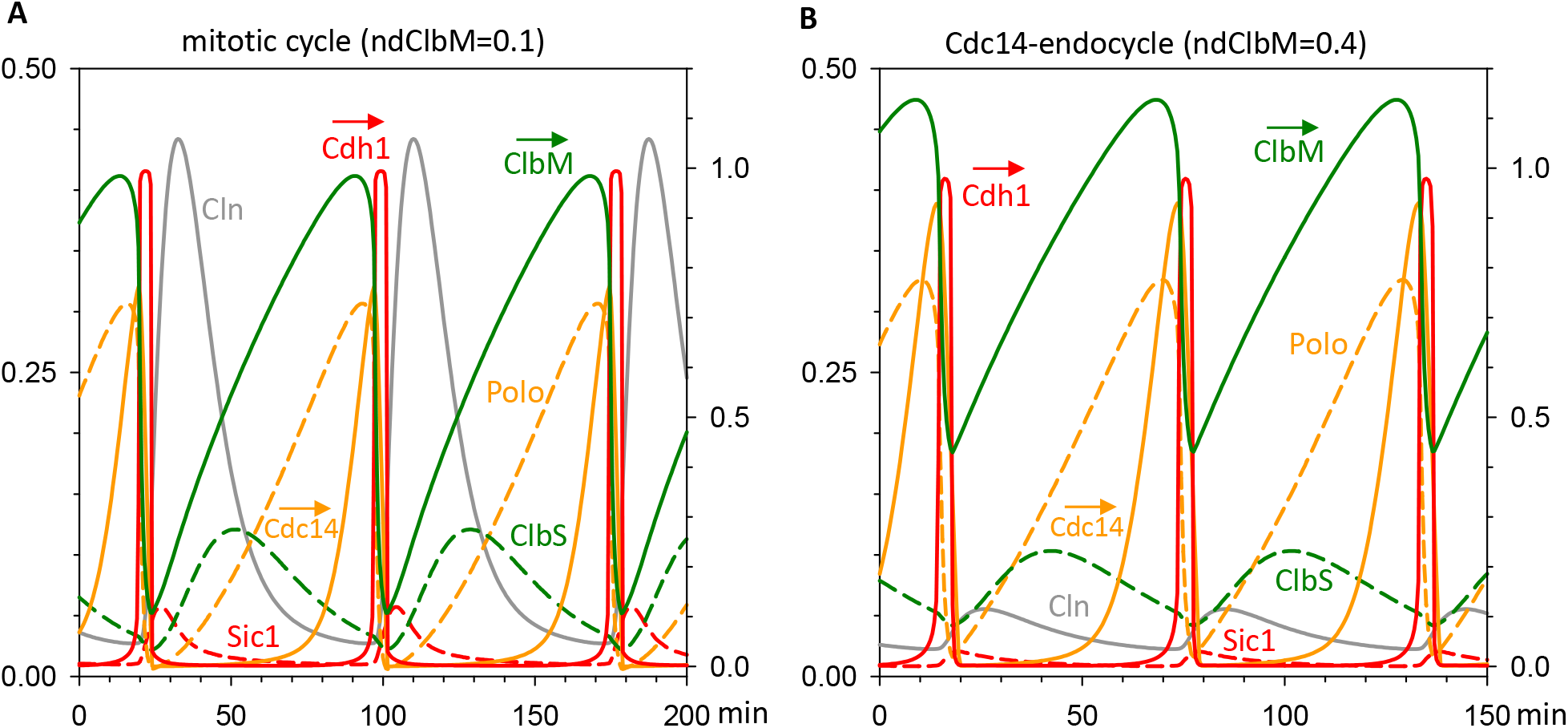
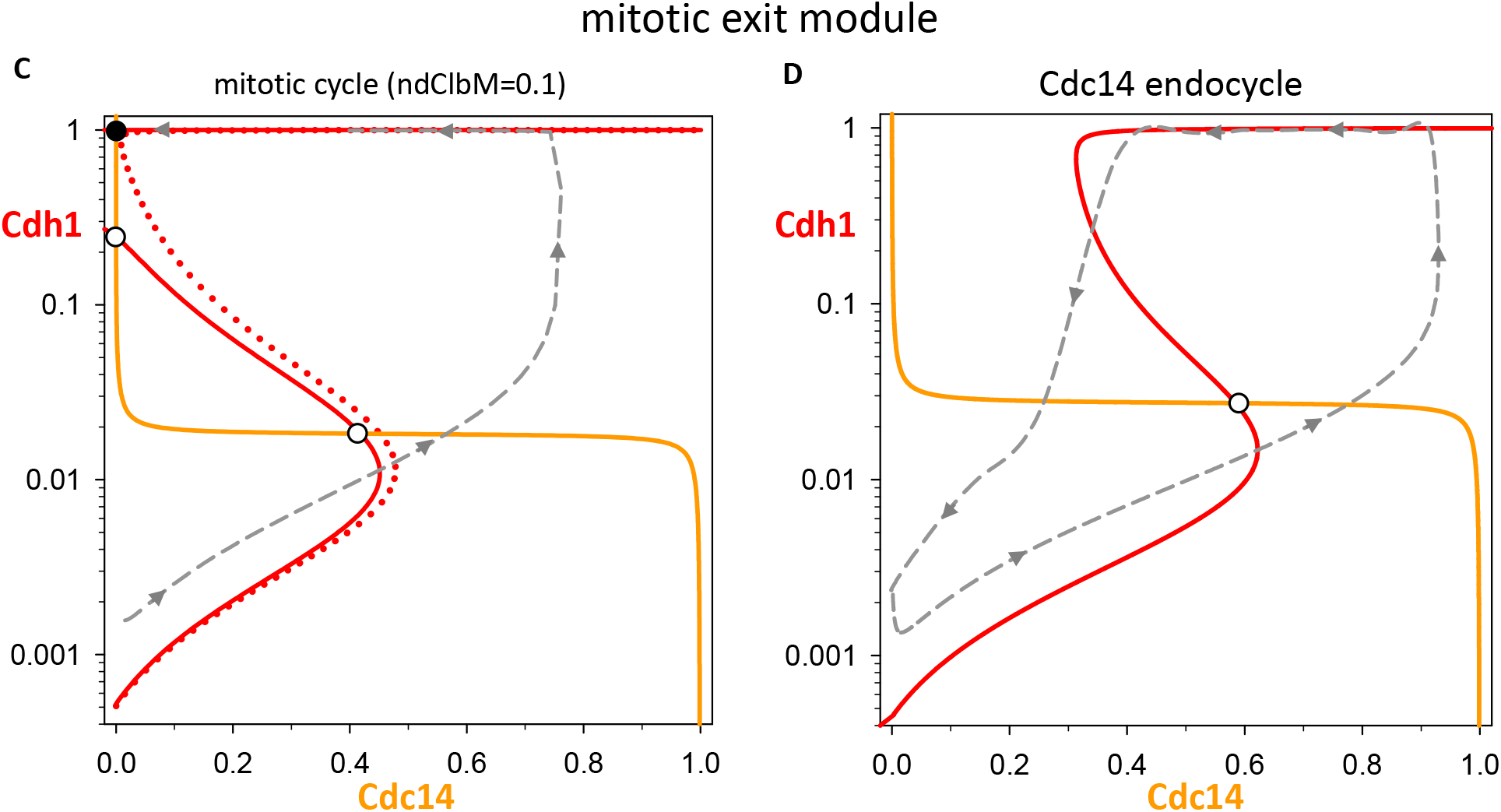
Cdc14 endocycles in the *CLB2kdΔ* mutant strain. **(A)** Numerical simulation of the model ODEs for ndClbM = 0.1. In this case, the cell executes normal mitotic cycles. **(B)** Numerical simulation for ndClbM = 0.4. In this case, the cell exhibits periodic activation and inactivation of Cdc14, without DNA synthesis or mitosis. **(C)** Pseudo-phase plane, for ndClbM = 0.1. Red curve is steady state activity of Cdh1 as a function of Cdc14; blue curve is steady state activity of Cdc14 as a function of Cdh1. Grey dashed line is a portion of the trajectory in panel A, projected onto the Cdc14-Cdh1 plane. The dotted red curve is drawn for ndClbM = 0.15, which is the SNIC bifurcation point. **(D)** For ndClbM = 0.4, the S-shaped (red) pseudo-nullcline has pulled into the positive quadrant of the pseudo-phase plane. The unique steady state (○) is unstable and surrounded by a limit cycle oscillation (grey dashed curve, derived from the simulation in panel B), which corresponds to Cdc14 endocycles. Although the dynamical system has six variables and the red and orange curves are not nullclines, the pseudo-phase plane gives a good idea of how Cdc14 endocycles arise from an inhibitor-amplified negative feedback loop in the presence of intermediate levels of non-degradable ClbM.

The existence of Cdc14 endocycles (Figure 5B) over a restricted range of expression of ndClbM is a consequence of the effect of non-degradable ClbM on the Cdh1 balance curve in Figure 3B (left side). For ndClbM = 0, the threshold for Cdh1 <u>inactivation</u> is at a <u>negative</u> value of Cdc14 (Figure 5C is a mirror image of Figure 3B). By increasing ndClbM, the Cdh1 inactivation-threshold moves to larger values of Cdc14, eventually crossing the y-axis at ndClbM ≈ 0.15 (the dotted red curve in Figure 5C). At this point the stable steady state (• on the solid red curve) collides with the unstable saddle point (○) underneath, and they both disappear. As a consequence, Cdh1 inactivation is no longer dependent on entering a new cell cycle and accumulating Cln- and ClbS-kinases; rather, the remnant ndClbM-kinase can inactivate Cdh1. In this case, there is only a single, unstable steady state of the control system, surrounded by a limit cycle oscillation (Figure 5D). (Suppl. Figure S4A shows that this limit cycle is independent of Cln and ClbS activities.) In the parlance of dynamical systems theory, this scenario is a SNIC bifurcation (‘saddle node on an invariant circle’). The limit cycle (the dashed curve in panel D) corresponds to the Cdc14 endocycles in panel B. The oscillation is a consequence of the fact that, as Cdc14 phosphatase activity drops, Cdh1 is inactivated by the remnant ClbM kinase activity in the cell, as evidenced by the Cdh1 inactivation-threshold at a positive value of Cdc14. The hysteretic nature of the oscillation is a consequence of the double-negative feedback loop between Cdh1 and the endogenous, degradable ClbM in the cell. It is an example of an ‘inhibitor-amplified negative feedback loop oscillator,’ in the terminology of Novak & Tyson [32].

Cdc14 endocycles persist over a limited range of expression of *CLB2kdΔ* (0.15 < ndClbM < 0.66), as observed by Lu & Cross [13], although our range is somewhat different from theirs. Because oscillations start at a SNIC bifurcation point, the period of oscillation drops significantly as ndClbM increases and the amplitude changes little (see Suppl. Figure 4B), as observed.

## Discussion

According to some molecular cell biology textbooks (see p. 870 of [33]) cell cycle transitions are rendered irreversible by proteolytic degradation of cell cycle regulators (like cyclins), because these processes are thermodynamically irreversible. In contrast, we have argued for many years [5, 15, 34] that irreversibility of cell cycle transitions is a consequence of bistable switches (‘latching gates’) created by systems-level positive and double-negative feedback motifs. The bistable switch is created by the fundamental opposition between cyclin B-dependent protein kinases (CDKs) and their antagonistic proteins: stoichiometric CDK inhibitors (CKIs), and E3 ubiquitin ligases (UbLs) that mediate degradation of S- and M-phase cyclins. The two stable steady states are G_1_ (antagonists active and CDKs inactive) and S-G_2_-M (*vice versa*). Unidirectional progression through the cell cycle is driven by ‘helper’ proteins: transcription factors that drive the G_1_ → S-G_2_-M transition, and ‘exit proteins’ that drive the reverse transition (from metaphase back to G_1_). Helper protein activities rise from near-zero to induce the transition and then fall back to low levels after the transition, because they are governed by negative feedback loops. The generic picture is illustrated in Figures 6A and B. In budding yeast cells, as we have explained, the transcription factors are MBF and SBF (Figure 6C) and the exit proteins are Cdc14 and Cdc20 (Figure 6D). It is the very robust switching of this latching gate between low and high CDK activity that drives the strict alternation of a proliferating cell lineage between G_1_ phase (unreplicated chromosomes) and S-G_2_-M phase (chromosome replication and partitioning in mitosis). <u>If the latches do not engage properly, as we have shown, then cells undergo ‘endocycles.’</u> (1) If mitotic cyclins are deleted, then a cell may undergo repeated rounds of DNA replication without mitosis (‘endoreplication cycles’), see Figure 4B. (2) If mitotic cyclins are not totally degraded in telophase, then a cell may undergo repeated cycles of exit-protein activation and inactivation (‘Cdc14 endocycles’ in budding yeast), see Figure 5D.

**Figure 6.**
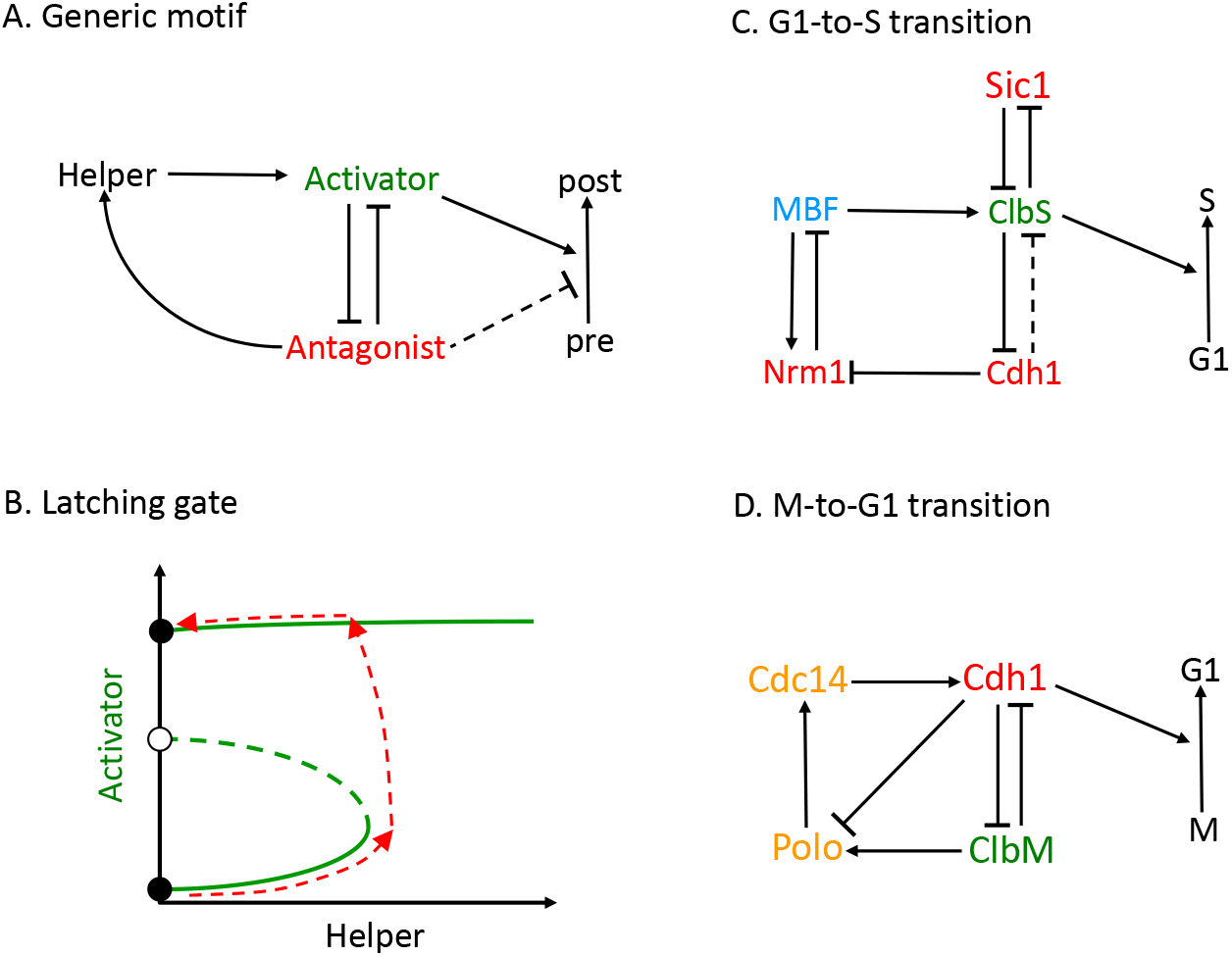
A general framework for cell-cycle transitions driven by bistable latching gates. **(A)** Generic motif. A bistable switch is created by the double-negative feedback between the ‘Activator’ of a cell-cycle transition and its ‘Antagonist’. The Antagonist may inhibit the transition. **(B)** Latching gate. Before the transition, the Antagonist dominates over the Activator (lower stable steady state •). To induce the transition, a ‘Helper’ protein rises from near-zero level (red dashed curve pointing right) and assists the Activator in overcoming the Antagonist (red dashed curve pointing up). Because the Helper is reliant on the Antagonist, Helper activity falls back to near-zero (red dashed curve pointing left) as the Antagonist is inactivated. **(C)** The G_1_-to-S transition in budding yeast. The Activator is ClbS-dependent kinase (Cdk1:Clb6) and the Helper is the transcription factor, MBF. The Antagonists are Cdh1 and Sic1 (Cdh1-mediated degradation of Clb6 is an assumption of our model). Cdh1 activates MBF by degrading its inhibitor, Nrm1. **(D)** The M-to-G_1_ transition in budding yeast. Notice that the Activator of this transition is Cdh1 and the Antagonist is ClbM (Cdk1:Clb1-4); the Helper is the protein phosphatase Cdc14. Polo kinase mediates the feedback signals from ClbM and Cdh1 to Cdc14. If the ‘latches’ fail to engage, then cells execute repeated rounds of DNA replication (see Figure 4B) or of Cdc14 sequestration and release (Figure 5D).

The reason behind each of these endocycles is the negative feedback signal that governs the Helper (either MBF or Cdc14). During the normal cell cycle, the Helper pushes the gate open, then the negative feedback signal pulls the gate closed and the latch catches the gate at a stable steady state (either G_1_ or M). In our view, endocycles are inadvertent consequences of this ‘logic’ of cell cycle control, when one component (the latching mechanism) fails. Proper sequencing of cell cycle events requires that the Helper proteins be inactivated after they have served their purpose (opening the gate). The negative feedback loop is analogous to the spring on a garden gate that pulls the gate closed behind the foot-traveler. If the latch doesn’t catch, then the gate will swing back and forth on the spring.

The generic motif in Figure 6A is one of the ‘Class 3’ motifs in Novak & Tyson’s classification of biochemical oscillators (see Figure 5c of [32]). The Class 3 schemes consist of ‘incoherently-amplified negative feedback loops’. In our case here, the negative feedback loop, Antagonist → Helper → Activator −| Antagonist, is ‘amplified’ by the double-negative loop between Activator and Antagonist, which is one leg of an incoherent feedforward loop: Antagonist <u>directly</u> inhibits Activator but <u>indirectly</u> activates Activator through Helper. As we stressed in [32], this motif is prone to sustained oscillations given sufficiently nonlinear interactions and a suitable selection of rate constant values. Normally the oscillations are suppressed by the latching property of the ClbM/Cdh1 interactions, but, if ClbM levels are constrained, then the Class 3 motif can oscillate. In our case here, there are two distinct Class 3 motifs (Figure 6C and D), which account for two different types of endocycles: ClbS endocycles (endoreplication) when ClbM level is constrained low, and Cdc14 endocycles when ClbM level is constrained high. In both cases, the oscillations arise via a SNIC bifurcation, which is characterized by finite amplitude and increasing frequency of oscillation as ClbM level moves past the bifurcation point.

Glycolytic oscillations in yeast cells are an example of this sort of inadvertent cycling, created by feedback signals required by the ‘logic’ of sugar metabolism…not by any apparent advantage of oscillations for the yeast cell. Phosphofructokinase (PFK), the first enzyme specific to six-carbon sugar catabolism, is allosterically inhibited by the ‘endproduct’ of the pathway (ATP). As ATP concentration increases, the flux of sugar through PFK decreases, conversely, if ATP concentration were to drop, then the flux would increase to replenish the ATP level. This regulatory effect is amplified by allosteric activation of PFK by ADP, whose concentration rises and falls in opposition to ATP. Because ATP is a substrate of PFK and ADP a product, these allosteric effects (necessary for proper regulation of energy metabolism) create a positive feedback on PFK activity, which, over a range of glucose input rates, generates sustained oscillations of carbon flux through PFK. (In the Novak-Tyson [32] classification, these are Class 2: inhibitor-amplified negative feedback loops.) The oscillations are tolerated because they seem to be of no particular advantage or disadvantage to the cells.

In a similar way, the logic of cell cycle control creates the possibility of endocycles under very specific circumstances. But, unlike harmless glycolytic oscillations, endoreplication cycles are deleterious for the cell, because they produce polyploid cells with 2^n^ copies of the genome and (worse yet) 2^n^ centrosomes, which makes subsequent sorting out of complete genomes nearly impossible. In yeast, polyploid cells nearly always die. Multicellular animals and plants have polyploid tissues, but they are terminally differentiated cell types that do not divide. Cdc14 endocycles, on the other hand, do not seem to be deleterious to yeast cells. Lu & Cross [13] showed that *GAL1-URL-CDC5* cells (which produce Cdc5 constitutively; i.e., they synthesize Cdc5 in G_1_ phase when ClbM-kinase activity is low) sometimes exhibit a second cycle of Cdc14 release from and reuptake by the nucleolus before re-entering the S-G_2_-M sequence quite normally.

Periodic cell-cycle events uncoupled from the CDK/Cdh1 ‘master’ control system are observed in some other circumstances. For example, Haase & Reed [35] observed periodic budding in a temperature-sensitive mutant strain *cdc4-3* of budding yeast, a strain that is defective in degrading Cln-type cyclins and Sic1, a stoichiometric inhibitor of Clb1-6. This mutant strain (at the restrictive temperature) fails to replicate its DNA or to divide, but undergoes periodic budding in the presence of high levels of *CLN2* mRNA and Cln2-dependent kinase activity. By expressing *CLN2* constitutively from a GAL promoter, the authors showed that fluctuations of mRNA level and kinase activity (above the high background levels) are not causative of periodic budding. So it is doubtful that periodic budding in the *cdc4-3* strain is a genuine ‘endocycle’ (i.e, an autonomous oscillator). In our opinion, periodic budding is likely driven by a constant high level of Cln2-kinase activity which drives the continuous accumulation of bud precursors, with buds appearing at regular intervals of time as bud primordia are assembled, as in the ‘structural model’ of [36] or the ‘dripping faucet’ model of Tyson et al. [37]. A similar mechanism seems to account for the formation of multiple septa in the *cdc16*^*ts*^ mutant strain of fission yeast [38]. Cdc16 is an inhibitor of the septation initiation network in fission yeast, so the mutant cells seem to have a constantly active SIN, which drives repeated septation events; see Figure 4 of [39]. For spindle pole body (SPB) reduplication in mutant strains of budding yeast, a similar story holds. SPB duplication requires both Clb5-6- and Cln1-2-dependent kinase activities and reduplication is strongly inhibited by Clb1-4-dependent kinases [40, 41]. SPB are periodically reduplicated in *clb1-5Δ* and *clb1-4Δ* strains [40].

Furthermore, strains that constitutively over-express high levels of Cln2 or Clb5 exhibit enhanced SPB reduplication [40], indicating that reduplication is not dependent on periodic oscillations of either Cln2 or Clb5. As the case for periodic budding, SPB reduplication is more likely a ‘dripping faucet’ than an autonomous limit cycle oscillator. This view is supported by the fact that centrosomes undergo repeated duplication in *Xenopus* embryos in the absence of either a nucleus or protein synthesis, and centrosome reduplication in frog egg extracts arrested in S phase is driven by the continuous presence of cyclin E [42].

In our opinion, periodic budding and SPB duplication in budding yeast that lack mitotic cyclins are not autonomous limit cycle oscillators, but periodic DNA replication and Cdc14 endocycles, as described here, are autonomous oscillators that arise when the latching mechanisms of cell-cycle control fail. Our view is consistent with the proposal of Manzoni et al. [14] that ‘events <u>capable</u> of repeating themselves multiple times [like Cdc14 endocycles or endoreplication cycles] are <u>restrained</u> to occur once per cycle by their coupling to the cyclin-Cdk engine’ (emphasis ours). It differs from the paradigm suggested by Lu & Cross [13] that ‘Cdc14 release, and likely other cell-cycle processes, are controlled by <u>intrinsically oscillatory</u> modules, that are <u>entrained</u> to a single occurrence at appropriate cell-cycle positions by cyclin-Cdk cycles through a “phase-locking” mechanism’ (emphasis ours). We do not view ClbS activation and Cdc14 release as <u>intrinsically</u> oscillatory modules that happen to be entrained by a complementary ClbM oscillator in a phase-locked relationship. Rather, we view the endocycles as inadvertent and deleterious oscillations that are normally <u>suppressed</u> by the CDK/Cdh1 latching-gate mechanism.

## Abbreviations

APC: anaphase promoting complex
CDK: cyclin-dependent kinase
CKI: cyclin-dependent kinase inhibitor
EP: exit protein
RP: restriction point
SAC: spindle assembly checkpoint
TF: transcription factor
UbL: ubiquitin ligase

