## Supplementary Material for "The oscillation of mitotic kinase governs cell cycle latches"

### Supplementary Text

#### *A simple model of the budding yeast cell cycle*

Our mathematical model provides a description of the *cdc20Δ pds1Δ clb5Δ* triple-mutant, in which Securin (Pds1), one of the S-phase cyclins (Clb5) and an APC/C activator (Cdc20) are deleted. We assume that the level of Cdc28 (Cdk1) is not limiting and the rate of Cdk1:Cyclin dimer accumulation is driven by the synthesis of different cyclins. The various Cdk1:Cyclin complexes are labelled by their cyclin component. The stoichiometric inhibitor, Sic1, binds to B-type cyclin dimers (Cdk1:Clb) to form inactive trimers. To distinguish these forms, we introduce the notation: *ClbM* = [Cdk1:ClbM] = 'free ClbM', $ClbM_t$ = [Cdk1:ClbM] + [Cdk1:ClbM:Sic1] = 'free ClbM' + 'complex with Sic1', and similarly for *ClbS* and $ClbS_t$. Furthermore, *Clb* = 'free ClbS' + 'free ClbM', and $Clb_t = ClbS_t + ClbM_t + ndClbM$, where $Clb_t$ refers to the total Clb concentration, including non-degradable ClbM (*ndClbM*) in some simulations. To keep the model simple, we assume that Sic1 binds reversibly to both S-phase and M-phase Cdk1 complexes with the same equilibrium dissociation constant ($K_{diss}$). In this case, the level of inactive Sic1:Clb complexes is given by:

$$Sic1Clb = \frac{2 \cdot Sic1_t \cdot Clb_t}{B + \sqrt{B^2 - 4 \cdot Sic1_t \cdot Clb_t}}$$

where $B = Sic1_t + Clb_t + K_{diss}$. The overall Clb activity (*ClbS* + *ClbM*) is calculated by subtracting the inactive complexes with Sic1 (*Sic1Clb*) from total Clb ($Clb_t$):

$$Clb = Clb_t - Sic1Clb.$$

Since SBF is inhibited only by the mitotic Clb, *ClbM* needs to be calculated as a fraction of Clb's not complexed by Sic1:

$$ClbM = \frac{ClbM_t + ndClbM}{Clb_t}(Clb_t - Sic1Clb)$$

The synthesis of Cln's (Cln1 & Cln2) is dependent on the SBF transcription factor, which is regulated by Cln activity (see below):

$$\frac{dCln}{dt} = k_{scln} \cdot SBF - k_{dcln} \cdot Cln$$

The synthesis of S-phase cyclin ClbS (Clb6) is driven by the MBF transcription factor, which is activated by Cln-phosphorylation and inhibited by Nrm1, a transcriptional repressor synthesized by MBF itself. The degradation of both ClbS and Nrm1 is controlled by APC/C in complex with Cdh1:

$$\frac{dClbS_t}{dt} = k_{sclbs} \cdot MBF_a - (k'_{dclbs} + k_{dclbs} \cdot Cdh1) \cdot ClbS_t$$

$$\frac{dMBF}{dt} = k'_{dis} \cdot (MBF_{tot} - MBF) - k'_{ass} \cdot MBF \cdot \left(Nrm1_t - (MBF_{tot} - MBF)\right)$$

$$\frac{dNrm1_t}{dt} = k_{snrm1} \cdot MBF_a - k_{dnrm1} \cdot Cdh1 \cdot Nrm1_t$$

In these equations, $Nrm1_t$ = *Nrm1* + *Nrm1MBF* = 'free Nrm1' + 'complex with MBF', $MBF_{tot}$ = *MBF* + *Nrm1MBF*, and $MBF_a = \frac{MBF \cdot Cln}{J_{MBF} + Cln}$ = 'active MBF', i.e., the fraction of free MBF that is phosphorylated by Cln (assuming rapid phosphorylation and dephosphorylation of MBF).

The synthesis of M-phase cyclins (Clb1-4) is autocatalytic [1] and their degradation is APC/C:Cdh1-dependent:

$$\frac{dClbM_t}{dt} = k'_{sclbm} + \frac{k_{sclbm} \cdot ClbM^n}{{J_{SBF}}^n + ClbM^n} - (k'_{dclbm} + k_{dclbm} \cdot Cdh1) \cdot ClbM_t$$

The synthesis of Polo-kinase is dependent on the activity of mitotic Cdk1 complexes, and it is degraded in a Cdh1-dependent fashion:

$$\frac{dPolo}{dt} = k_{spolo} \cdot ClbM - \left(k'_{dpolo} + k_{dpolo} \cdot Cdh1\right) \cdot Polo$$

We assume that Sic1 is synthesized at a constant rate and that its degradation is controlled by dual phosphorylation [2]. The first phosphorylation step, which is assumed to be in equilibrium with dephosphorylation, is catalysed by both Cln- and Clb-kinases, while the second step is Clb-dependent:

$$\frac{dSic1_t}{dt} = k_{ssic} - \left[k'_{dsic} + k_{dsic} \cdot Clb \cdot \frac{Cln + Clb}{J_{sic1} + Cln + Clb}\right] \cdot Sic1_t$$

The switch-like activation and inactivation of three regulatory proteins (SBF, Cdh1 and Cdc14) is described by 'Goldbeter-Koshland kinetics'. Cln-kinases phosphorylate and inactivate the inhibitor (Whi5) of the SBF transcription factor, i.e., SBF is activated by Cdk1:Cln, and SBF activity is inhibited by mitotic Cdk1 (ClbM, [1]):

$$\frac{dSBF}{dt} = \frac{\left(k'_{asbf} + k_{asbf} \cdot Cln\right) \cdot (1 - SBF)}{J_{SBF} + 1 - SBF} - \frac{k_{isbf} \cdot ClbM \cdot SBF}{J_{SBF} + SBF}$$

Cdh1 is inactivated by both S-phase and M-phase Cdk1 (recall, *Clb* = *ClbS* + *ClbM*) and reactivated by Cdc14 phosphatase:

$$\frac{dCdh1}{dt} = \frac{(k'_{acdh1} + k_{acdh1} \cdot Cdc14) \cdot (1 - Cdh1)}{J_{cdh1} + 1 - Cdh1} - \frac{(k'_{icdh1} \cdot Cln + k_{icdh1} \cdot Clb) \cdot Cdh1}{J_{cdh1} + Cdh1}$$

The nucleolar release of Cdc14 is induced by Polo-kinase:

$$\frac{dCdc14}{dt} = \frac{k_{acdc14} \cdot Polo \cdot (1 - Cdc14)}{J_{cdc14} + 1 - Cdc14} - \frac{k_{icdc14} \cdot Cdc14}{J_{cdc14} + Cdc14}$$

This set of ordinary differential equations is implemented in the subsequent ‘ode’ file for simulation with XPPAUT (http://www.math.pitt.edu/~bard/xpp/installonwindows.html). The ‘basal’ parameter values for all simulations are given in the following table.

**Parameter values for simulations:**

| |
|---|
| **Cln synthesis/degradation & SBF regulation:** |
| $k_{scln} = 0.2$, $k_{dcln} = 0.2$, $k'_{asbf} = 1$, $k_{asbf} = 10$, $k_{isbf} = 25$, $J_{sbf} = 1$ |
| **ClbS synthesis and degradation:** |
| $k_{sclbs} = 0.15$, $k'_{dclbs} = 0.1$, $k_{dclbs} = 0.05$ |
| **MBF regulation by Nrm1:** |
| $k_{snrm1} = 0.05$, $k_{dnrm1} = 0.1$, $MBF_{tot} = 0.5$, $k'_{ass}$=1, $k'_{diss}$=0.001, $J_{mbf} = 0.01$ |
| **ClbM synthesis and degradation:** |
| $k'_{sclbm} = 0.01$, $k_{sclbm} = 0.01$, $k'_{dclbm} = 0.01$, $k_{dclbm} = 1$, $J_{clbm} = 0.01$, $n = 2$ |
| **Polo synthesis/degradation & Cdc14 regulation:** |
| $k_{spolo} = 0.01$, $k'_{dpolo} = 0.01$, $k_{dpolo} = 1$, $k_{acdc14} = 1$, $k_{icdc14} = 0.25$, $J_{cdc14} = 0.01$ |
| **Sic1 synthesis/degradation & Clb binding :** |
| $k_{ssic} = 0.02$, $k'_{dsic} = 0.01$, $k_{dsic} = 2$, $J_{sic1} = 0.01$ $K_{diss} = 0.05$ |
| **Cdh1 activation & inactivation:** |
| $k'_{acdh1} = 1$, $k_{acdh1} = 10$, $k'_{icdh1} = 0.2$, $k_{icdh1} = 10$, $J_{cdh1} = 0.01$ |
| **Nondegradable ClbM:** |
| ndClbM=0 for mitotic and endoreplication cycle and ndclbM >0 for Cdc14 endocycle. |

**XPPAut ode code for simulation**

```
# XPPAut model for cell cycle latches
# Differential equations
Cln' = kscln*SBF - kdcln*Cln
ClbSt' = ksclbs*MBFa - (kdclbs' + kdclbs*Cdh1)*ClbSt
MBF' = kdiss'*(MBFtot - MBF) - kass'*MBF*(Nrm1t - (MBFtot - MBF))
Nrm1t' = ksnrm1*MBFa - kdnrm1*Cdh1*Nrm1t
ClbMt' = ksclbm' + ksclbm*ClbM^n/(Jclbm^n + ClbM^n) - (kdclbm' + kdclbm*Cdh1)*ClbMt
Polo' = kspolo*ClbM - (kdpolo' + kdpolo*Cdh1)*Polo
Sic1t' = kssic' - (kdsic' + kdsic*Clb*(Cln+Clb)/(Jsic1+Cln+Clb))*Sic1t
SBF' = (kasbf' + kasbf*Cln)*(1-SBF)/(Jsbf + 1 - SBF) - kisbf*ClbM*SBF/(Jsbf + SBF)
Cdh1' = (kacdh1' + kacdh1*Cdc14)*(1 - Cdh1)/(Jcdh1 + 1 - Cdh1) - (kicdh1'*Cln +
kicdh1*Clb)*Cdh1/(Jcdh1 + Cdh1)
Cdc14' = kacdc14*Polo*(1 - Cdc14)/(Jcdc14 + 1 - Cdc14) - kicdc14*Cdc14/(Jcdc14 + Cdc14)

# Algebraic equations for Clb & Sic1 binding
MBFa = MBF*Cln/(Jmbf + Cln)
Clbt = ClbSt + ClbMt + ndClbM
BB = Sic1t + Clbt + Kdiss
Sic1Clb = 2*Sic1t*Clbt/(BB + sqrt(BB^2 -4*Sic1t*Clbt))
Clb = Clbt - Sic1Clb
ClbM = (ClbMt+ndClbM)*(Clbt - Sic1Clb)/Clbt

# Auxiliary variables
aux ClbS = ClbSt*(Clbt - Sic1Clb)/Clbt
aux ClbM = (ClbMt+ndClbM)*(Clbt - Sic1Clb)/Clbt

# Parameter values
p kscln=0.2, kdcln=0.2, kasbf'=1, kasbf=10, kisbf=25, Jsbf=1
p ksclbs=0.15, kdclbs'=0.1, kdclbs=0.05
p ksnrm1=0.05, kdnrm1=0.1, MBFtot=0.5, kass'=1, kdiss'=0.001, Jmbf=0.01
p ksclbm'=0.01, ksclbm=0.01, kdclbm'=0.01, kdclbm=1, Jclbm=0.05, n=2
p kspolo=0.01, kdpolo'=0.01, kdpolo=1, kacdc14=1, kicdc14=0.25, Jcdc14=0.01
p kssic'=0.02, kdsic'=0.01, kdsic=2, Jsic1=0.01, Kdiss=0.05
p kacdh1'=1, kacdh1=10, kicdh1'=0.2, kicdh1=10, Jcdh1=0.01
p ndClbM=0

# XPP instructions
@ METH=stiff, XLO=0, XHI=250, YLO=0, YHI=1, total=250, dt=0.25, XP=time
@ NPLOT=8,YP=Cln,YP2=Cdh1,YP3=Sic1t,YP4=Nrm1t,YP5=Polo,YP6=Cdc14,YP7=ClbS,YP8=ClbM
done
```

### Supplementary Figures

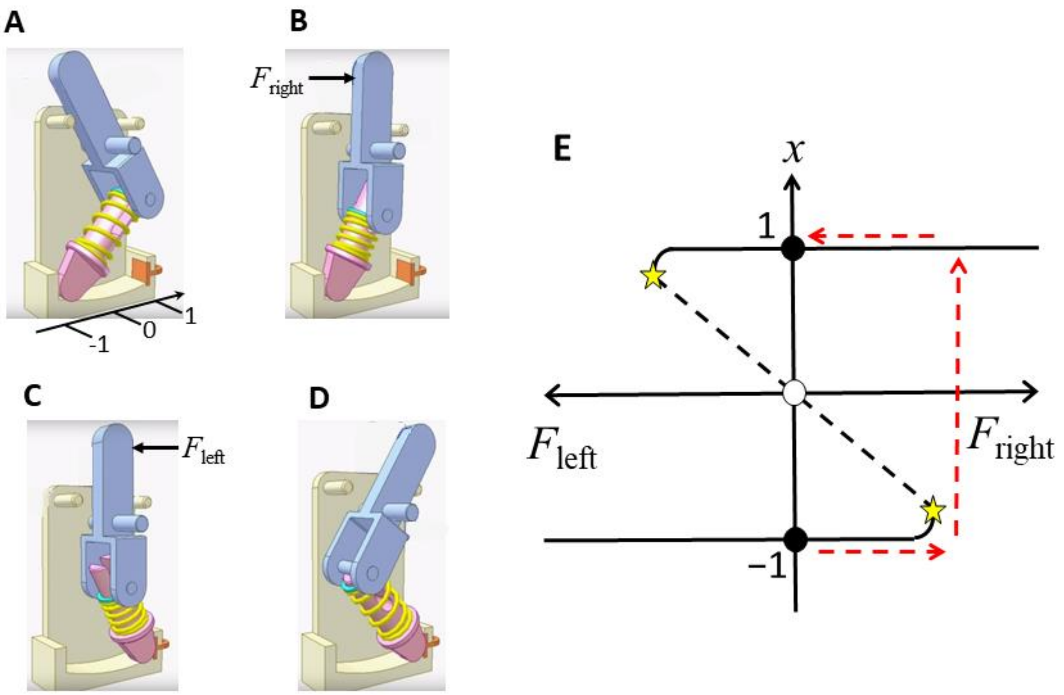


**Suppl. Figure S1.** A mechanical latching switch. **(A)** The switch is in the off position; the spring is relaxed. **(B)** A rightward force applied to the lever depresses the spring but not enough to flip the switch. **(C)** If pushed a little harder to the right, the compressed spring will snap back to its longer (lower energy) state, and the purple headpiece will press against the orange contact, closing an electrical circuit. In this state, there is no applied force. **(D)** Now a leftward force depresses the spring but not enough to flip the switch. If pushed a little harder to the left, the spring will snap the switch to the off position (A). **(E)** Position ($x$) of the pink head piece, measured in distance from the middle of the slide (we assume a frictionless slide), as a function of force on the lever. When $F = 0$, the headpiece can rest at either the left stop or the right stop ($x = -1$ or 1), as denoted by the stable steady state symbol ●. For $F = 0$, there is also an unstable steady state (○) where the hinged piece is straight up-and-down and the spring is maximally compressed. Any small perturbation to the right or left will allow the spring to snap the headpiece to one side or the other. To flip the switch on (the red dashed lines), the user must apply a rightward force large enough to align the two hinged pieces (maximal compression of the spring at the yellow star). Beyond that point, the spring snaps the headpiece into contact with the orange terminal. The force on the lever drops abruptly to zero as it pulls away from the user's finger. To turn off the switch (not shown), the user must apply a leftward force to the lever in the same manner.

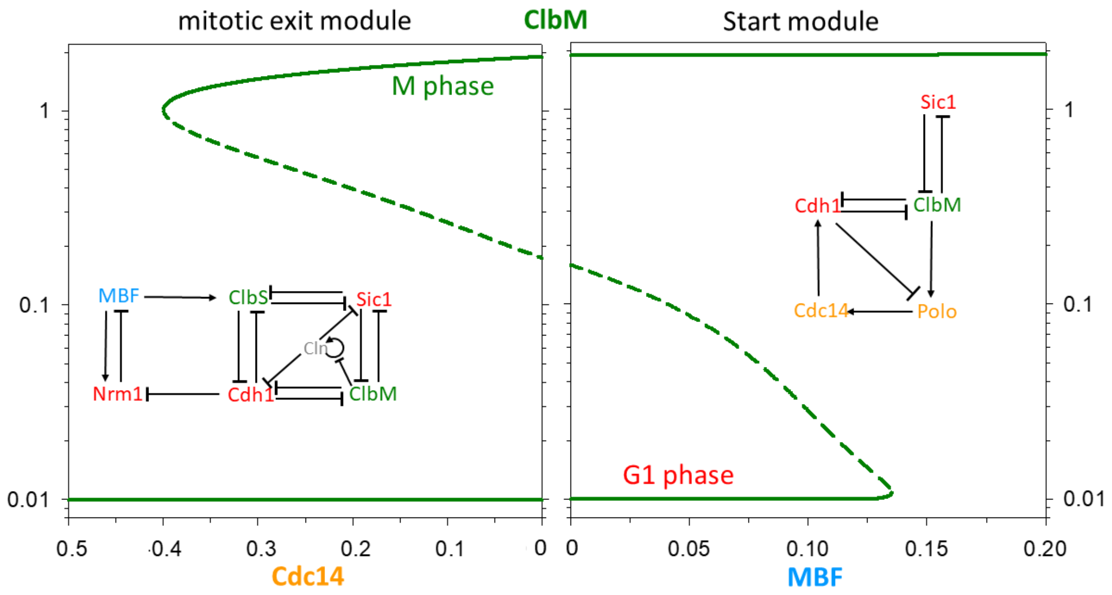


**Suppl. Figure S2.** Alternative view of the latching diagram for budding yeast. Same as Figure 3 in the text, except we plot steady state level of ClbM in dependence on MBF and Cdc14. When MBF = Cdc14 = 0, the control system is bistable: $G_1$ phase with ClbM concentration very low (0.01) and Cdh1 activity high (1 in text Figure 3); M phase with ClbM concentration high (2) and Cdh1 activity low (0.0005); and an unstable steady state with intermediate values of ClbM (0.15) and Cdh1 (0.1). This figure confirms the schematic illustration of bistability and hysteresis in Figure 9 of Chen et al. [3].

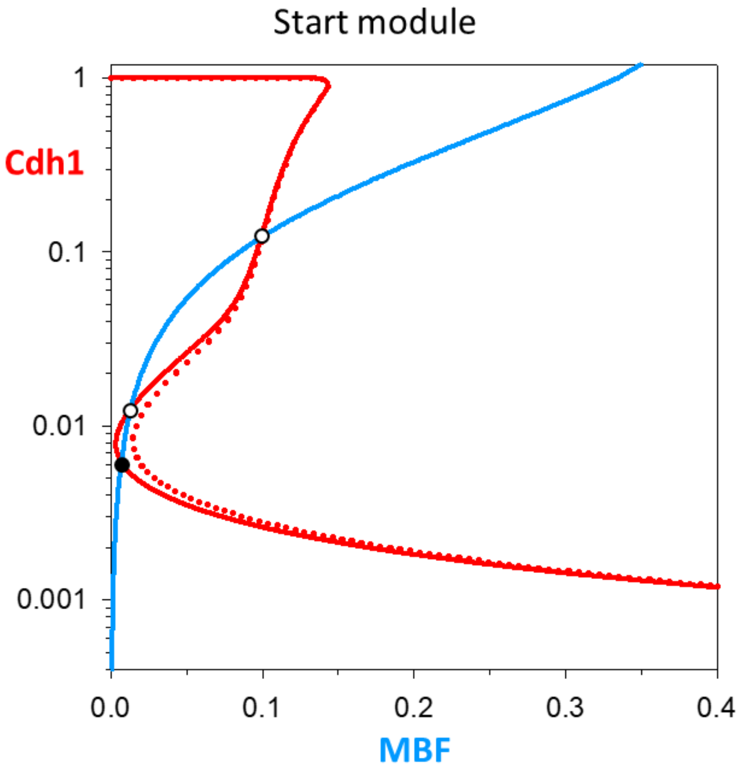


**Suppl. Figure S3.** Endoreplication cycles arise as limit cycle oscillations from a SNIC (saddle-node on an invariant circle) bifurcation. We draw the pseudo-phase plane for $k_{sclbm}$'= $k_{sclbm}$=0.00225, which lies between their values of 0.01 for normal cycling, Figure 3 (right panel), and their values of zero for endoreplication cycles, Figure 4B. The red nullcline (Cdh1 at steady state) has moved to the right, so that the saddle point (○) and the node (●) are very close together. For parameter values slightly smaller (0.0021; dotted red curve), the saddle and node have coalesced and disappeared, leaving only a stable limit cycle of very long period around the unstable node. The defining characteristic of SNIC-derived limit cycles is that the period is initially very long and drops rapidly as the bifurcation parameter pulls away from the bifurcation point (see Figure S4B).

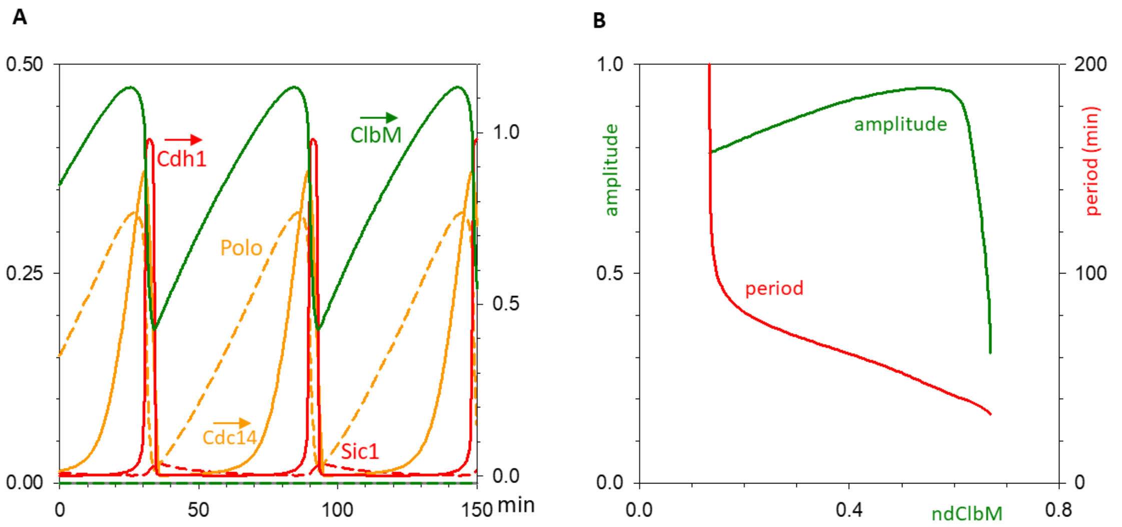


**Suppl. Figure S4.** Cdc14 endocycles. **(A)** In the absence of Cln and ClbS ($k_{scln} = k_{sclbs} = 0$), simulated Cdc14 endocycles are nearly identical to Figure 5B. **(B)** Bifurcation diagram. We plot the period (red) and amplitude (green) of Cdc14 endocycles as functions of non-degradable ClbM. The oscillations arise at ndClbM = 0.15 by a SNIC (saddle-node on an invariant circle) bifurcation and are extinguished at ndClbM = 0.66 by a SNPO (saddle-node of periodic orbits) bifurcation. Notice the abrupt drop of period as ndClbM increases beyond the SNIC bifurcation point, which agrees with the observations of Lu & Cross [4].
